# Direct In-Sample Sequencing of the 3′ Transcriptome Expands the Capabilities of Optical Pooled Screens

**DOI:** 10.1101/2025.10.11.681797

**Authors:** Daniel Honigfort, Pedro Belda-Ferre, David White, Kousik Sundararajan, Mariam Dawood, Jake LeVieux, Jacob Moreno, Xiaodong Qi, Kyle Metcalfe, Duuluu Naranbat, Andrew Altomare, Connor Thompson, Carlos Ruiz Perez, Bryan Lajoie, Hyukjin J. Kwon, Pazinah Bhadha, Tristin Rammel, John Rabalais, Matthew Kellinger, Semyon Kruglyak, Sinan Arslan, Michael Previte

**Affiliations:** Element Biosciences, San Diego, CA

## Abstract

We present a platform that directly sequences single guide RNAs and endogenous 3′UTRs in fixed cells while simultaneously measuring protein abundance and cellular morphology. We demonstrate platform capability by performing optical pooled screening of CRISPR-perturbed lung cancer cells. This approach unites direct in-sample RNA sequencing with complementary phenotypic readouts, enabling comprehensive, scalable, and functional genomics analyses within a single experiment.

## Main

A major challenge in genomics is linking gene perturbations with cellular phenotypes, particularly when responses to perturbations or treatments are transient and occur at varied time points. Pooled CRISPR screening has transformed functional genomics by enabling large-scale, systematic interrogation of gene function, but most current assays typically capture only a single phenotypic layer at a time.^1,2^ Optical pooled screening (OPS)^3,4^ adds image-based readouts of cellular and organelle morphology, yet remains limited to relatively small sets of cell-painting markers.^5^ Conversely, Perturb-seq delivers transcriptome-wide, single-cell profiles but lacks direct measurements of protein abundance, morphology, or spatial context.^6^ The field therefore faces an unmet need: methods that can connect genetic perturbations to multimodal phenotypes across multiple time points in a pooled and scalable manner.

Our approach enables direct in-sample sequencing of single guide RNAs and endogenous 3′UTRs while simultaneously quantifying protein abundance and cellular morphology, providing a comprehensive view of gene expression and cellular response. We implemented these capabilities in Teton Atlas, an automated and scalable multiomic platform built on the AVITI24 system. The workflow proceeds as follows. After culture and fixation on the flowcell, each cell is probed with six cell-painting features (generating over 500 cell profiling features from Cell Profiler^7^), a 50-plex protein detection panel, OPS guide detection, and unbiased 3′ transcriptome profiling (**Fig. 1A**).

**Figure 1:**
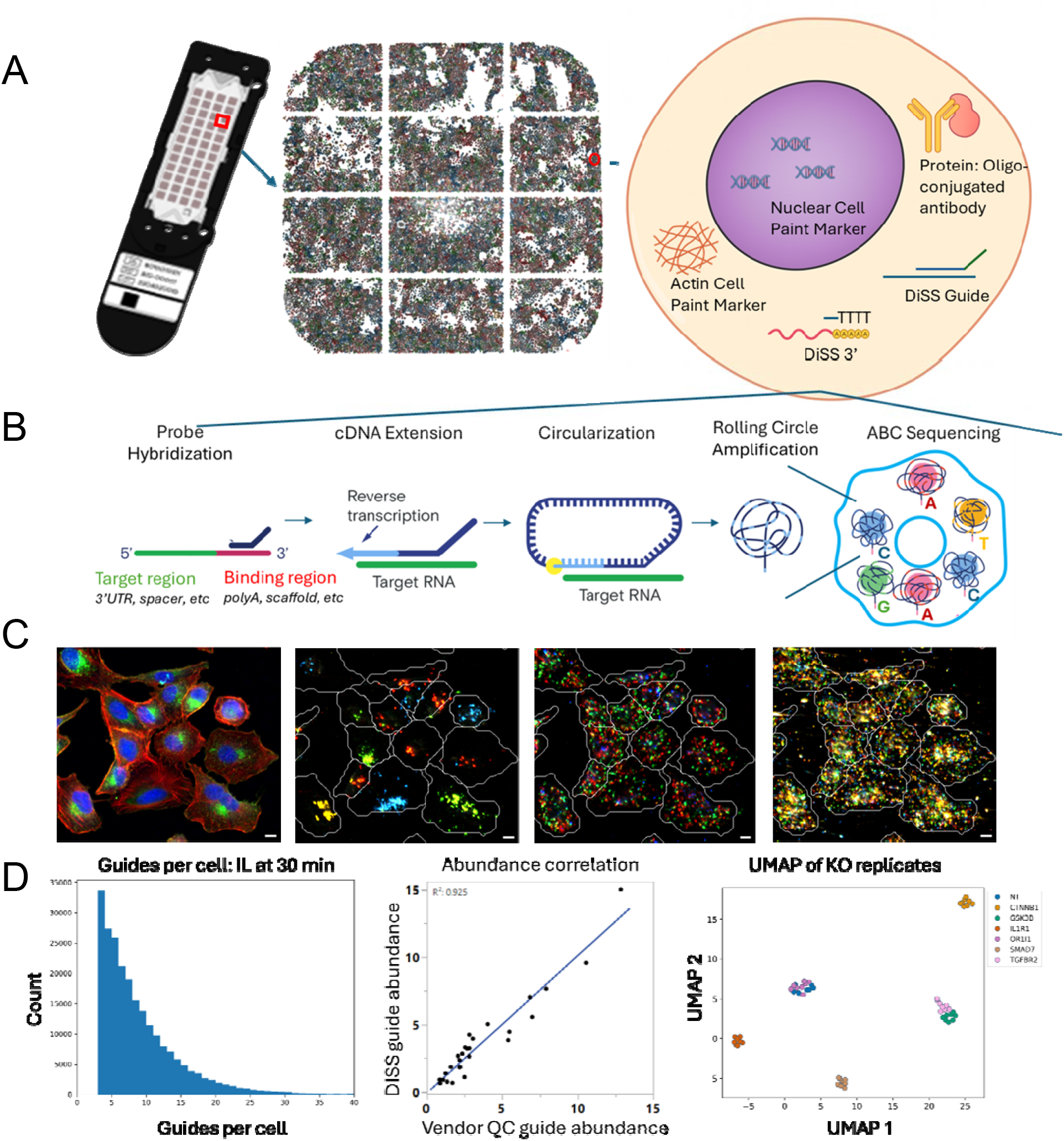
(A) Visual of the flowcell where cells are fixed and analyzed. Zoom into a single well to see the population of cells on the surface. Zoom into a cartoon of the assay being performed inside each cell on the AVITI24 instrument. (B) Overview of DISS chemistry, which is initiated by hybridization of a single sided DNA probe, which is then reverse transcribed and circularized before amplification by rolling circle amplification. (C) Images of the same cells with multiomic images superimposed over segmentation. 1. Cell Paint, with membrane (green) Nucleus (blue), and Actin (red) 2. sgRNA detection with DISS, 3. 3’ Transcriptome with DISS, 4. Protein detection. For basecalling assignment, A=red, C=blue, T=yellow, G=green. Scale bar is 20 μm. (D-F) QC measures including guide sequences per cell (D), DISS guide abundance correlation with 2D NGS (E), and UMAP of KO replicates across guides and wells (F). There are 8 points for each gene, corresponding to the 4 sgRNA targeting that gene in each of 2 replicate wells.

The central advance of this work is an improved method for direct optical sequencing of RNA molecules within intact cells^8–11^ termed Direct In-Sample Sequencing (DISS). DISS chemistry begins with the hybridization of a DNA probe oligonucleotide (**Fig. 1B**). For transcriptome profiling of 3’ UTRs, the DNA probe hybridizes specifically to the poly(A) tail of mRNA; for CRISPR guide detection, the DNA probe hybridizes to the invariant sgRNA scaffold 3’ to the variable spacer sequence. Next, reverse transcription extends the probe to encode the downstream RNA sequence as cDNA. The resulting probe-cDNA construct is ligated to form a circular DNA anchored to its endogenous RNA, preserving spatial information. Rolling Circular Amplification (RCA) is used to amplify the circular DNA into a concatemer and is sequenced using Avidite Based Chemistry (ABC).^12^ The entire process of DISS chemistry through ABC is automated on the fluidics based-platform AVITI24 system^13^, enabling robust, scalable operation with minimal user interaction. DISS is integrated with parallel measurement of high-plex protein abundance via the sequencing of oligo-conjugated antibodies, and morphological measurements derived from cell painting markers, as previously described (**Fig.1C**).^13,14^

A key feature of DISS chemistry is its use of single-sided DNA probes: short oligonucleotides that hybridize to a single known sequence adjacent to the target region and are then extended to capture downstream RNA. This design contrasts sharply with molecular inversion (“padlock”) probes commonly used in optically pooled CRISPR screens, which require prior knowledge of both 3′ and 5′ flanking sequences around a guide spacer. Such constraints often necessitate engineered expression systems, which fuse guides to synthetic mRNA elements to provide a constant upstream region.^15^ By requiring only a single hybridization site, DISS probes remove this design limitation and enable direct detection of unmodified sgRNAs expressed from standard CRISPR constructs. The same principle enables unbiased 3′ UTR profiling by designing probes to hybridize with the poly(A) tail of mRNAs. This streamlined architecture supports robust guide detection and broad transcriptome profiling without the need for engineered constructs or large probe panels.

For platform evaluation, we applied DISS to a pooled CRISPR screen in A549 lung cancer cells expressing Cas9. The 28-guide library included four guides each targeting CTNNB1, GSK3B, IL1R1, SMAD7, and TGFBR2, along with non-targeting (NT) control and guides to OR1I1, an olfactory receptor with no expected phenotype changes in this cell type. These guides induce perturbations in key genes in inflammation, Wnt, and TGF pathways. We selected this system because these pathways are well studied and provide both predictable benchmarks and diverse, measurable responses across molecular and cellular modalities. The effects of these CRISPR perturbations were quantified under standard growth conditions, as well as following stimulation with IL-1 or TGF ligands over a 48-hour time course. In the main text we focus on IL-1 and the first 4 time points, where protein, RNA, and morphology measurements converged most strikingly.

DISS successfully assigned over 80% of cells to a unique perturbation, with a median of six sgRNA reads per cell (**Fig. 1D**). Detected guide abundances correlated closely with measurements from orthogonal genomic sequencing, indicating minimal detection bias (**Fig. 1E**). Furthermore, transcript counts for four of the six targeted genes were significantly reduced relative to controls, consistent with knockout-induced nonsense-mediated decay (**Supplementary Fig. 1**); SMAD7 and OR1I1 were too low in abundance for reliable assessment. To assess the concordance between replicate wells and guides targeting the same gene, we performed UMAP analysis from the pseudo-bulk 3’ transcriptome profiles for each sgRNA across replicate wells (**Fig. 1F**). Perturbations targeting the same gene clustered closely together, while non-targeting and OR1I1 controls overlapped, demonstrating consistent transcriptome measurements and high concordance between replicates

To validate the platform’s ability to resolve pathway dynamics, we characterized the cellular response to IL-1β stimulation, a well-defined innate immune signaling pathway directly perturbed in our screen through knockout (KO) of the IL-1β receptor IL1R1.^16,17^ As expected, IL-1β induced robust protein signaling in control cells but not in IL1R1-KO cells. After 30 min of stimulation, multiple proteins were significantly differentially regulated (**Fig. 2A**), including rapid phosphorylation of p38 and its downstream substrate HSP27, followed by delayed accumulation of p53 (**Fig. 2B, Supplementary Fig. 2A-B**).^18,19^ These responses, associated with inflammation-linked survival and stress pathways, were markedly attenuated in IL1R1-KO cells, which remained near baseline. Importantly, unrelated OR1I1 knockouts behaved like non-targeting controls.

**Figure 2:**
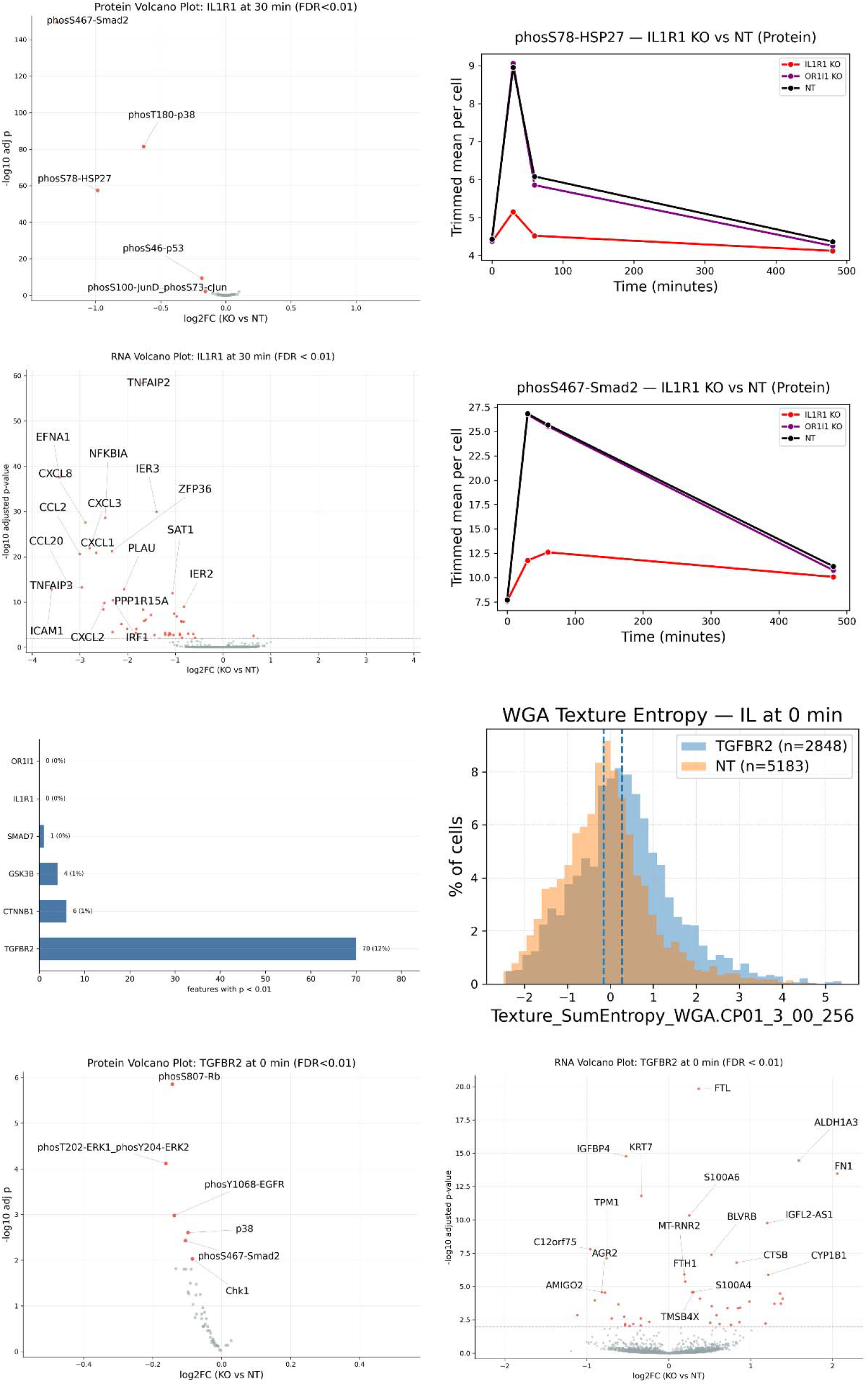
(A) Volcano plot of differential protein abundance (IL1R1-KO vs NT) at 30 min IL-1β; MAPK panel (50-plex). (B) Phospho-HSP27 over the IL-1β time course (0–4 h); early surge in NT, absent in IL1R1-KO. (C) Volcano plot of differential RNA abundance (IL1R1-KO vs NT) at 30 min IL-1β. (D) Phospho-SMAD2 over the IL-1β time course (0–4 h); strong response in NT, minimal in IL1R1-KO. (E) Count of Cell Painting features differing vs NT by KO; TGFBR2-KO shows markedly more changes. (F) Single-cell distributions for Texture_SumEntropy_WGA (WGA membrane marker); TGFBR2-KO shifted vs NT. (G) Volcano plot of differential protein abundance (TGFBR2-KO vs NT), untreated; MAPK panel (50-plex). (H) Volcano plot of differential RNA abundance (TGFBR2-KO vs NT) at time 0.

3′ Transcriptome profiling with DISS supported these findings. After 30 min of IL-1β treatment, IL1R1-KO cells showed broad attenuation of the transcriptional inflammatory response, including reduced expression of NF-κB-associated transcripts (CXCL8, CCL2, NFKBIA, ZFP36, JUN, EGR1) compared to controls (**Fig. 2C, Supplementary Figure 2C**).^20^ Temporal analysis of differentially expressed genes revealed distinct expression clusters reflecting the dynamic nature of inflammatory signaling (**Supplementary Fig. 3**), and gene ontology analysis highlighted enrichment of perturbed genes within the TNF, NF-κB, and IL-17 pathways (**Supplementary Fig. 4**).^21^ Together, these results confirm that IL1R1 is essential for initiating canonical IL-1 signaling and demonstrates the platform’s ability to capture coordinated protein and transcriptional responses to perturbation.

IL-1β signaling was not limited to canonical inflammatory pathways. We detected robust Smad2 phosphorylation following IL-1β stimulation, a result we did not anticipate but that aligns with prior observations in the literature^22,23^. Under IL-1β stimulation, phospho-Smad2 increased sharply in non-targeting controls but was abolished in IL1R1 knockouts, indicating IL-1–dependent crosstalk to Smad2 (**Fig. 2D**). Notably, this IL-1– induced Smad2 phosphorylation persisted in TGFBR2 knockouts, demonstrating that the crosstalk does not require canonical TGF receptor signaling (**Supplementary Fig. 5A**). By contrast, when cells were treated with TGFβ, Smad2 phosphorylation was eliminated by TGFBR2 knockout (**Supplementary Fig. 5B**), confirming canonical pathway dependence. Together, these findings place the IL-1–driven crosstalk upstream of Smad2 and downstream of IL1R1, but independent of TGFBR2-mediated signaling.

To assess how perturbations affect basal cell state in the absence of external stimuli, we examined steady-state phenotypes across our CRISPR panel. Morphological profiling revealed that TGFBR2 knockout had the most pronounced impact, with 12% of >500 CellProfiler features significantly altered relative to non-targeting controls (**Fig. 2E**). Single-cell feature distributions showed clear shifts in membrane and cytoskeletal organization (**Fig. 2F**), indicating broad effects on cellular architecture that align with known links between TGF signaling and cytoskeletal dynamics.^24,25^

To investigate the mechanistic basis of these changes, we analyzed protein and transcriptomic profiles under basal conditions. TGFBR2 knockout cells exhibited reduced phosphorylation of Smad2 and ERK proteins (**Fig. 2G**) and differential expression of numerous genes involved in cytoskeletal regulation (SEMA3C, ARHGEF2, TPM1), adhesion (FN1, CD44), and stress response (FTL, IGFBP4, BLVRB) (**Fig. 2H**). These complementary readouts link the observed morphological changes to underlying signaling and transcriptional programs, providing a mechanistic view of the effects of the knockout.

Together, these results demonstrate that loss of TGFBR2 reprograms the basal state of epithelial cells, altering homeostatic signaling, gene expression, and morphology. In contrast, OR1I1 knockouts showed no significant changes across any modality, confirming the specificity of the assay (**Supplementary Fig. 6)**.

## Discussion

This work demonstrates that integrating direct in-sample sequencing of guides and RNA with targeted protein and morphological measurements provides complementary layers of information for interpreting genotype–phenotype relationships in pooled CRISPR screens. Protein profiling captures rapid phosphorylation dynamics, while transcriptome readouts reveal broader network remodeling that shapes long-term cellular states. Together, these modalities resolve both the immediate signaling consequences of perturbation and the downstream transcriptional programs that sustain phenotypic change.

In the IL1R1 knockout, the combination of measurements revealed a collapse of canonical inflammatory signaling, and the absence of a direct transcriptional signature accompanying Smad2 phosphorylation highlighted the importance of protein-level measurements for detecting transient events. The observation that some of the largest effects were confined to the 30-60 minute time points further underscores the need for temporal resolution in functional screens. In the TGFBR2 knockout, integrating transcriptomic and proteomic profiles provided mechanistic insight into the substantial morphological remodeling observed at steady-state.

Together, these results highlight the platform’s strength in linking genetic perturbations to complex phenotypes across different timescales by integrating morphology, protein, and transcriptome measurements in a pooled, scalable assay. This integrated approach establishes a foundation for future screens that seek to map how perturbations propagate across multiple molecular layers and over time.

## Methods

### Cell culture

A549 cells stably expressing CAS9 and transfected with the 28 guide library were purchased as a custom library from Cellecta. Cells were cultured at 37 °C with 5% CO2 in F12K media (10-025-CV, Corning) supplemented with 10% (vol/vol) FBS (A3257207RP, Gibco), 1% (vol/vol) penicillin–streptomycin (15140-122, Gibco), 10 ug/ml Blasticidin (SBR00022, Sigma Aldrich), and 0.5 ug/ml Puromycin (AAJ67236XF, Fisher Scientific).

### Quantification of guide abundance

The relative abundance of cells expressing each given sgRNA of the library was performed using PCR and NGS of the sgRNA sequences from genomic DNA using the NGS Prep Kits for sgRNA shRNA and DNA barcode libraries (LNGS-101, Cellecta).

### Cytokine treatment

A549 cells with CRISPR edits were lifted with 0.25% Trypsin (25200-056, Gibco) and seeded directly onto Element 12-well slide kit (810-00012, Element Biosciences) at a density of 5k cells/well. Cells were allowed to adhere for 12hrs, then treatment was started in a staggered fashion such that they were all fixed with 4% Formaldehyde (F8775, Sigma) for 20min concurrently with the longest timepoint. Exogenous ligands were added directly to the growth media in the following concentrations: 5ng/mL TGFβ1 (H8541, Sigma Aldrich), 100ng/mL TNFa (T6674, Sigma Aldrich), 10ng/mL IL1b (IL038, Sigma Aldrich), 100ng/mL Wnt-3a (315-20, Fisher Scientific).

### Cytoprofiling Assay

Guide sequencing and protein detection were performed using an early R&D version of the Teton Atlas™ Cartridge and Reagent Kit - Low Output (PN: 860-00040) on the AVITI24 platform (Element Biosciences) with 3 batches of 3’ Transcriptome, 1 batch of protein, 1 batch of OPS, and 6 cell paint markers. Protein targets were selected from the Teton Human MAPK & Cell Cycle Panel Kit (Part No. 860-00040). Fixed flow cells were loaded onto the AVITI24, where both protein barcodes and DISS probes containing spacer cDNA were amplified, imaged, and sequenced using Avidite Based Chemistry (ABC). Image data were processed with a basecalling algorithm to decode barcodes or spacer sequences, which were spatially registered to individual cell boundaries for per-cell protein quantification and downstream analysis.

### Statistical analysis (protein and RNA)

All differential analyses were performed separately for each KO and timepoint, comparing KO to matched non-targeting (NT) controls. For both protein and RNA, per-cell counts were aggregated to guide×well samples and then collapsed across the two replicate wells within each timepoint, yielding four guide-level replicates per KO and four NT guide replicates. Each KO×timepoint set was analyzed with DESeq2 (via PyDESeq2) using a negative binomial GLM with design ~ condition (KO vs NT). DESeq2 returned log□ fold-changes, Wald p-values, and Benjamini–Hochberg FDR (padj).

### Volcano plots

Protein and RNA volcanoes were drawn from the DESeq2 results: x-axis = log2FC (KO vs NT), y-axis = −log10(padj when available, otherwise p), with significant points highlighted and top hits auto-labeled; figures were exported as PNGs alongside the source tables. When noted in the figure legends, highlighted points satisfied |log2FC| > 0.4 and FDR < 0.01.

### Morphology analysis and breadth plot

At each timepoint, morphology features were first aggregated to guide×well means. For each feature, KO (≈8) vs NT (≈8) was tested with a two-sided Mann–Whitney U (MWU) test (exact when possible, otherwise asymptotic); Benjamini–Hochberg correction was applied within each KO×time. We report p-values, FDR (padj), median differences (KO−NT), and rank-biserial correlations. For the breadth panel, we summarized, per KO and timepoint, the number and fraction of significant features (e.g., nominal p < 0.01 and/or FDR < 0.05) and plotted those counts as bars.

### Rationale for MWU on morphology (vs DESeq2)

Morphology measurements are continuous intensities/texture metrics, not counts; their distributions can be non-normal and feature-specific in scale. MWU is a non-parametric, rank-based test that remains valid under these conditions and with small n, making it well-suited for comparing KO vs NT guide×well summaries. In contrast, DESeq2’s NB model is tailored to count data (with library-size normalization), which does not apply directly to morphology features; hence MWU provides the appropriate inference here.

## Notes

### Competing Interest Statement

All authors are current or former employees of Element Biosciences and may hold stock options in the company

