## Supplementary Figures for "Direct In-Sample Sequencing of the 3′ Transcriptome Expands the Capabilities of Optical Pooled Screens"

Supplementary Information

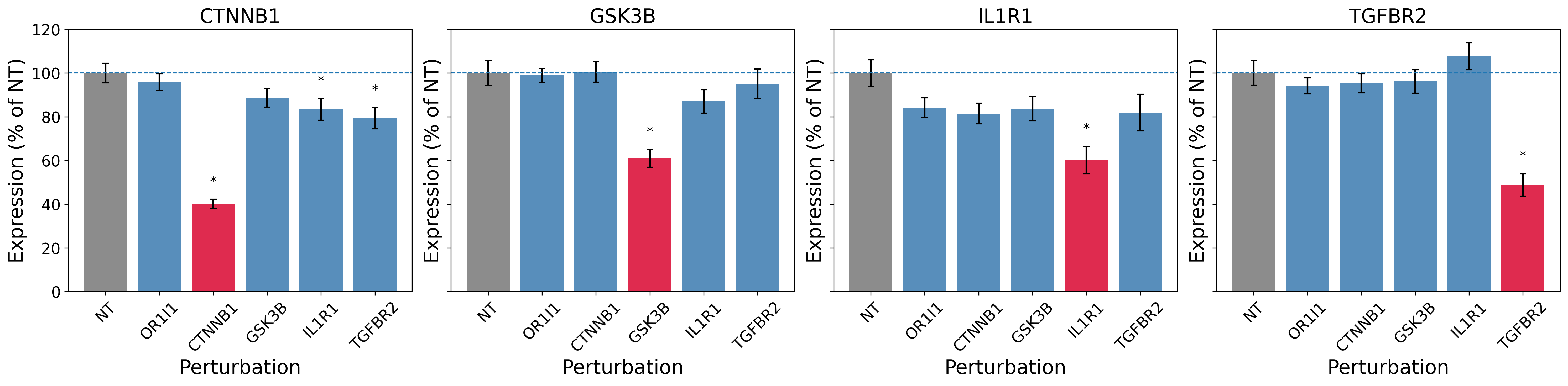

Supplementary figure 1: Suppression of transcripts associated with the corresponding KO. For each KO shown, only the corresponding transcript (red bar) is significantly reduced relative to the NT control as assessed via Mann-Whitney test. For each transcript, the height of the bar is the expression level as a percentage of the NT control (grey). The modest decrease in IL1R1 expression seems initially at odds with the large downstream impact of the KO. One possibility is that the altered transcript is not completely removed via nonsense mediated decay, despite the CRISPR edit. However, no functional protein can be formed, explaining the downstream impact on other transcripts and proteins.

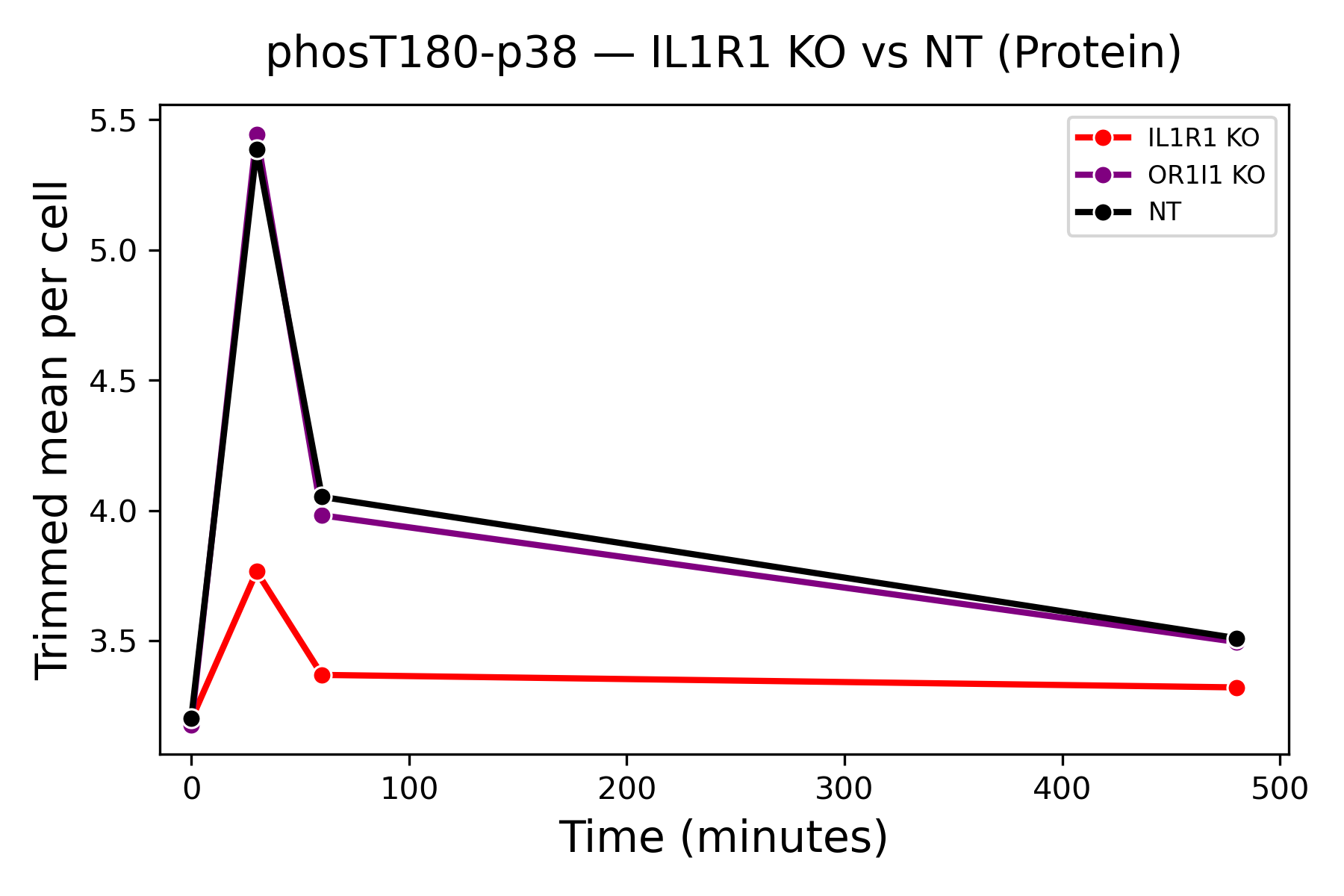

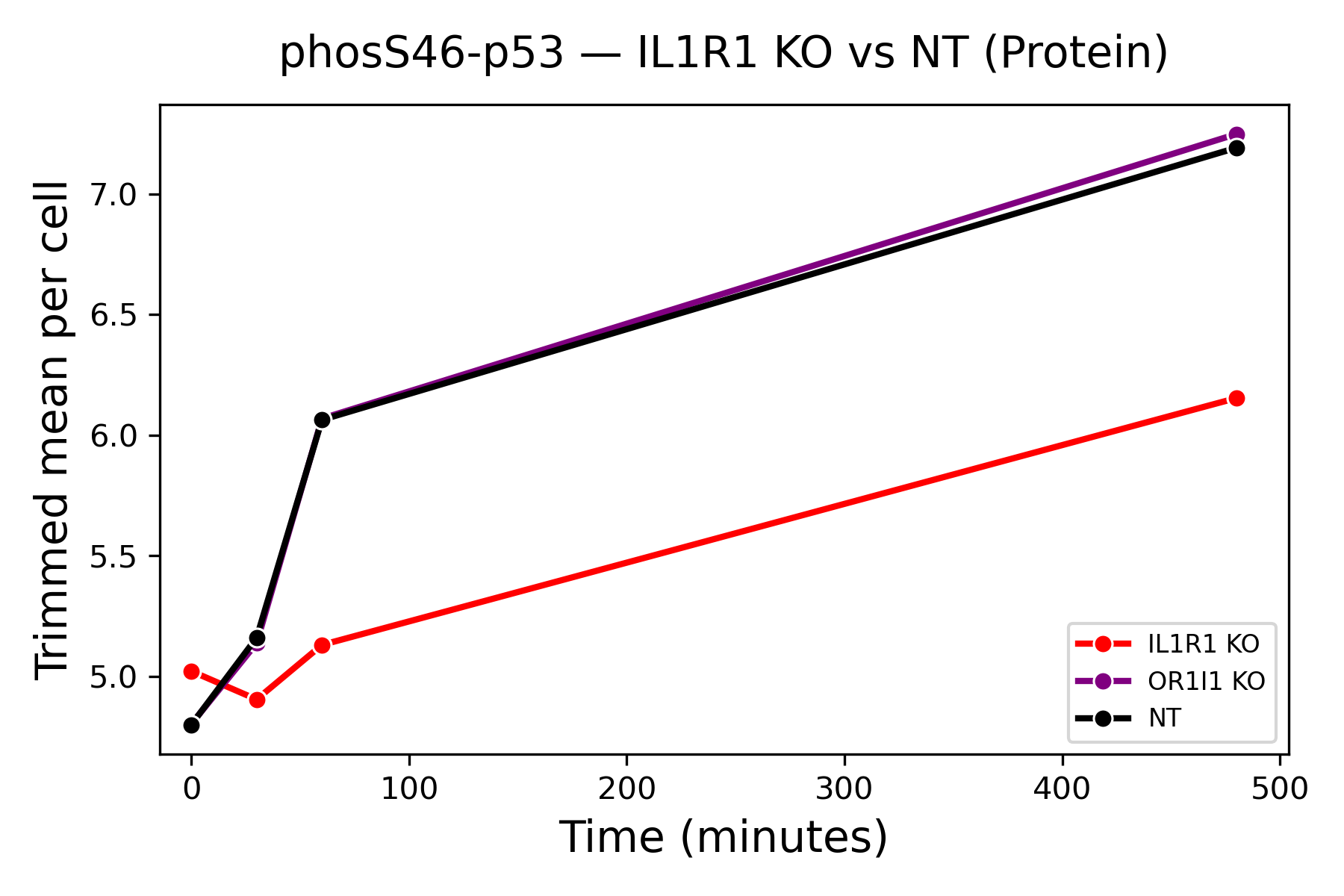

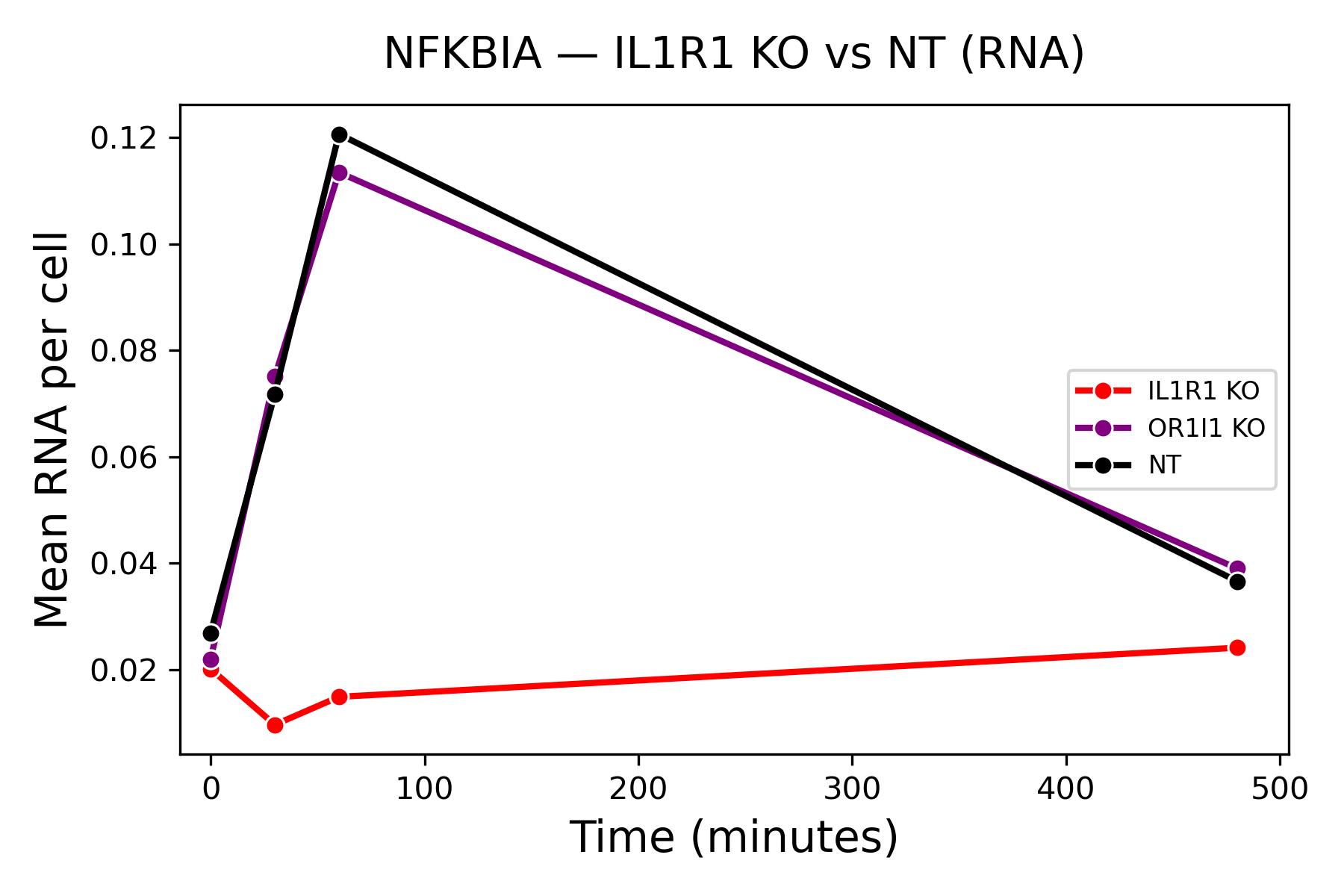

A

B

C

Supplementary figure 2: Impact of the IL1R1 KO on protein and mRNA targets. (A) phospho-p38 response in IL1R1 KO vs NT. (B) phospho-p53 delayed response in IL1R1 vs NT. (C) NFKB1A mRNA response in IL1R1 vs NT. Note that the OR1I1 perturbation behaves similarly to the NT control as expected.

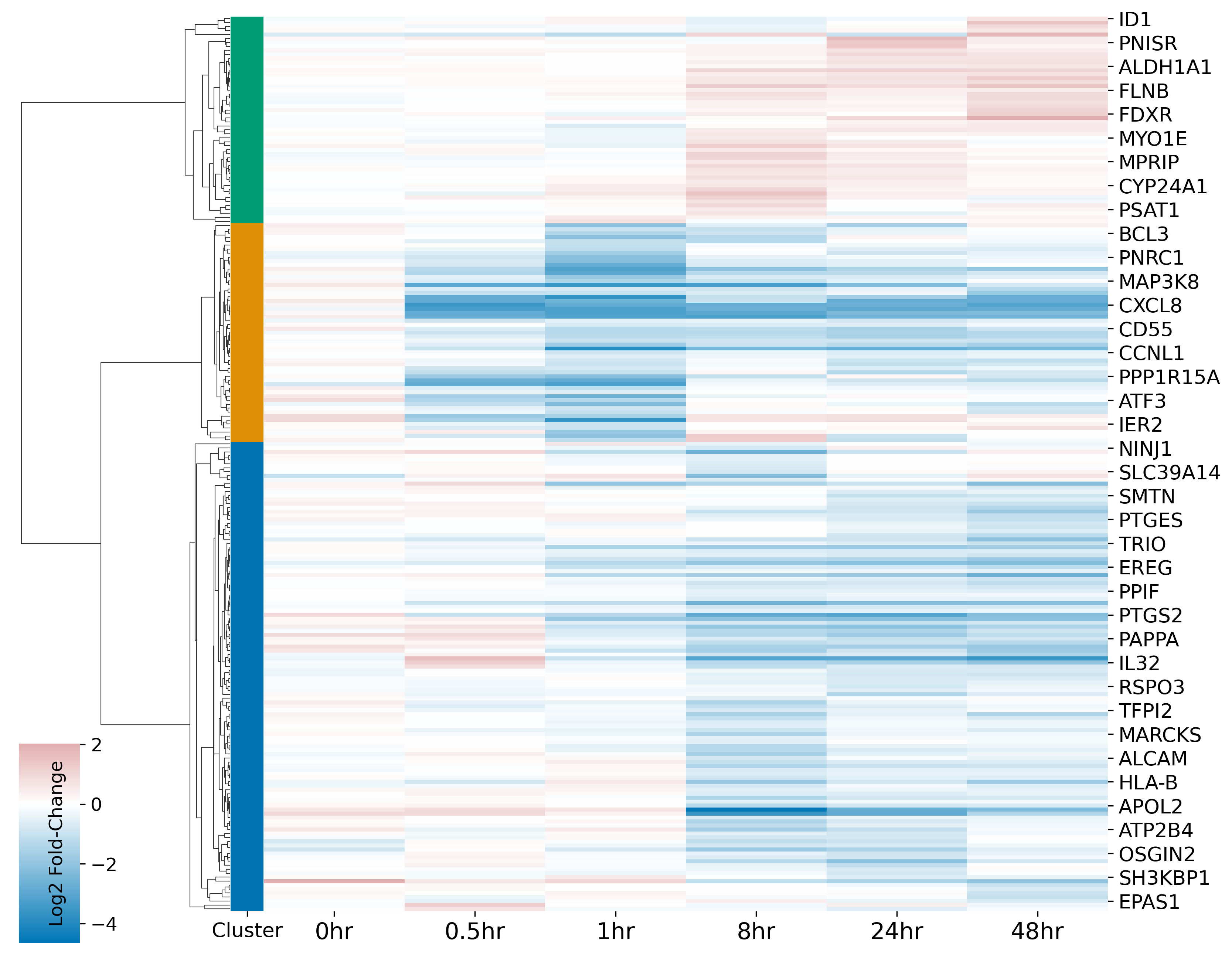

Supplementary figure 3: Heatmap of log₂ fold-change values for differentially expressed (DE) genes between IL1R1 KO and NT control cells identified over time. Each row represents a gene identified as DE at least once, and each column corresponds to a treatment time point. Genes were hierarchically clustered using cosine distances and average linkage based on their temporal expression changes relative to NT control cells. K-nearest neighbor (KNN, k = 10) clustering was applied to group genes with similar expression dynamics; three clusters were specified to facilitate visualization and comparison of major expression patterns. Row colors denote the KNN-derived gene groups.

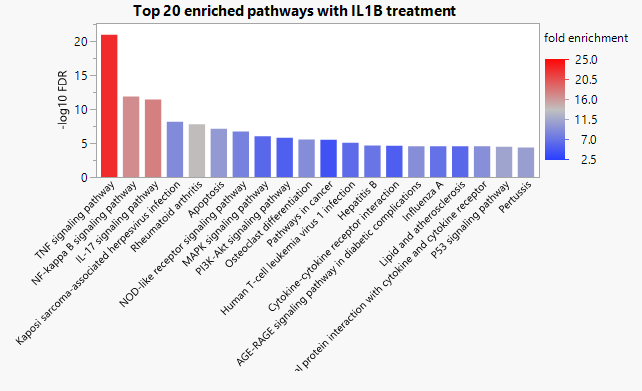

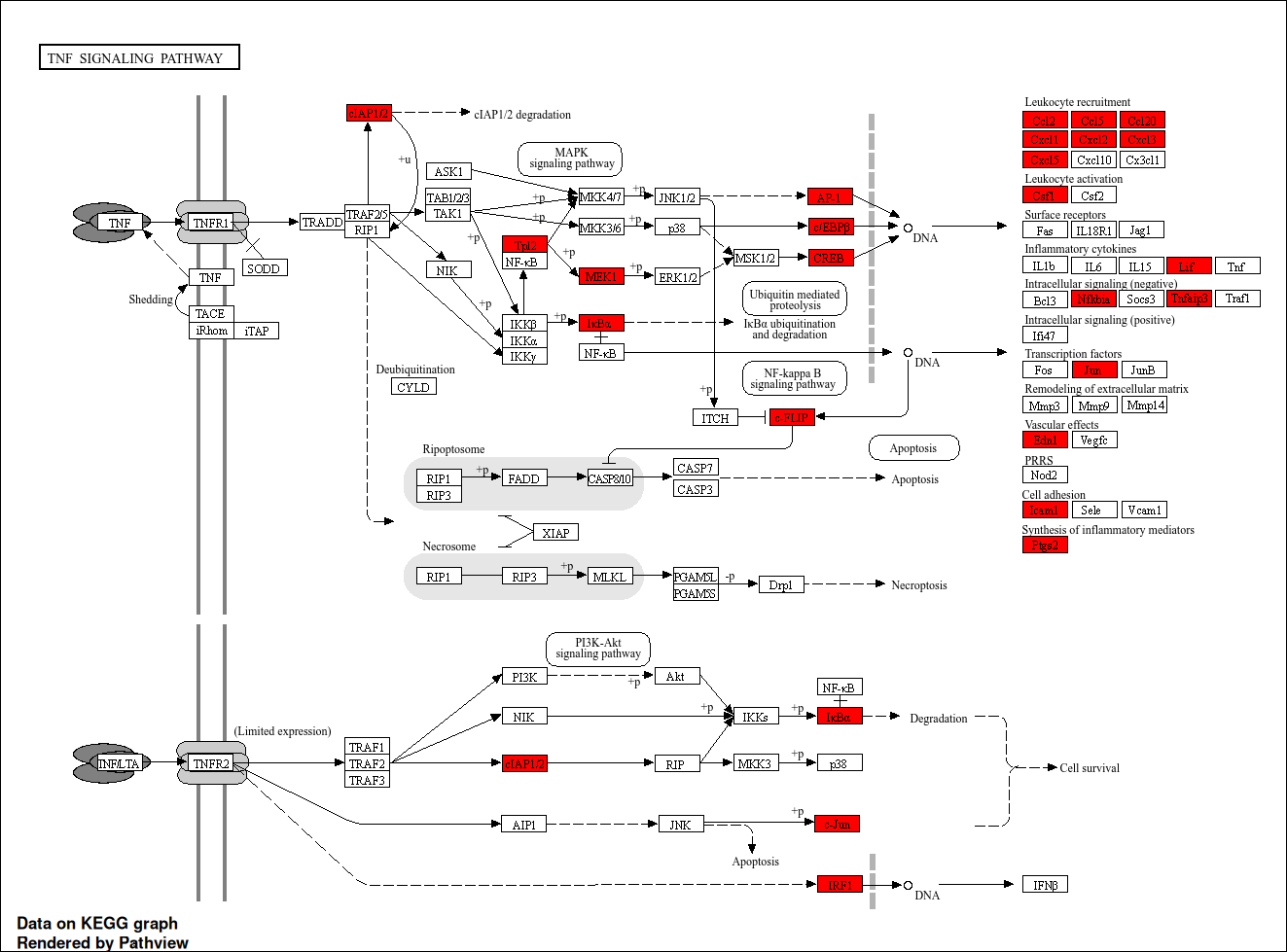

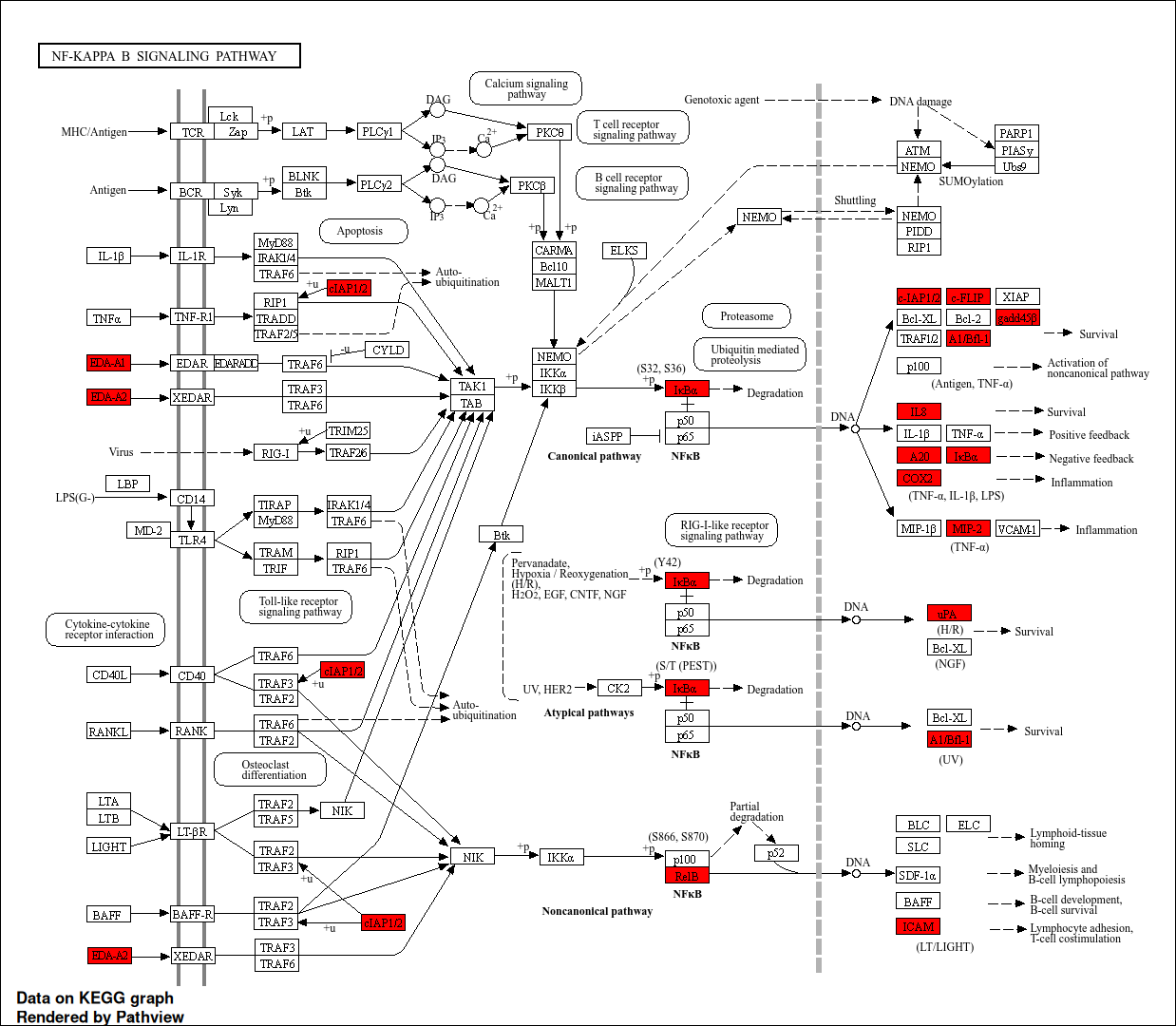

Supplementary figure 4: Gene Ontology analysis from ShinyGO of significant differentially regulated genes between IL1R1 knockout and NT control when treated with IL1B. (A) top 20 enriched KEGG pathways by fold-enrichment. (B) the most enriched KEGG pathway for TNF inflammatory signaling. TNF is a similar pro-inflammatory cytokine to the IL1β used in this study. (C) The second most enriched KEGG pathway for differentially regulated genes was NF-kappa B signaling. NF-κB pathway is a central regulator of inflammation. Genes highlighted in red were significantly differentially regulated between NT and IL1R1 knockout cells during IL1β treatment.

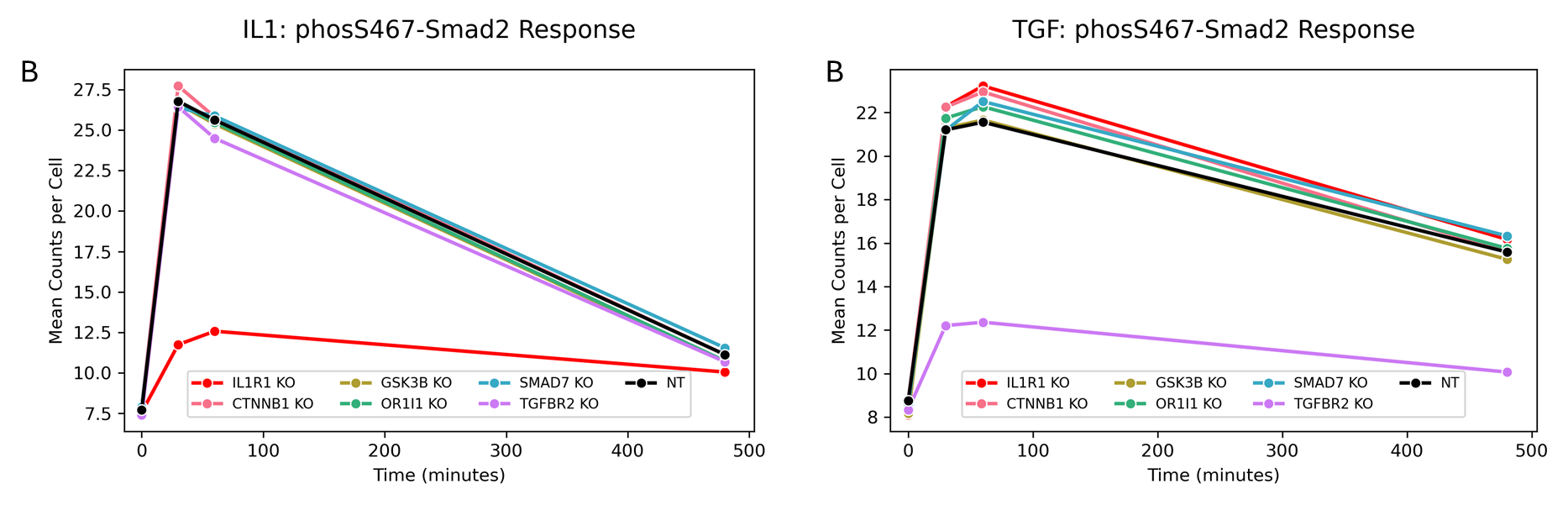

Supplementary figure 5: Phospho-Smad2 response across KOs present in CRISPR pooled library. (A) IL1B treatment timecourse from 0 to 480 min. (B) TGF treatment timecourse from 0 to 480 min.

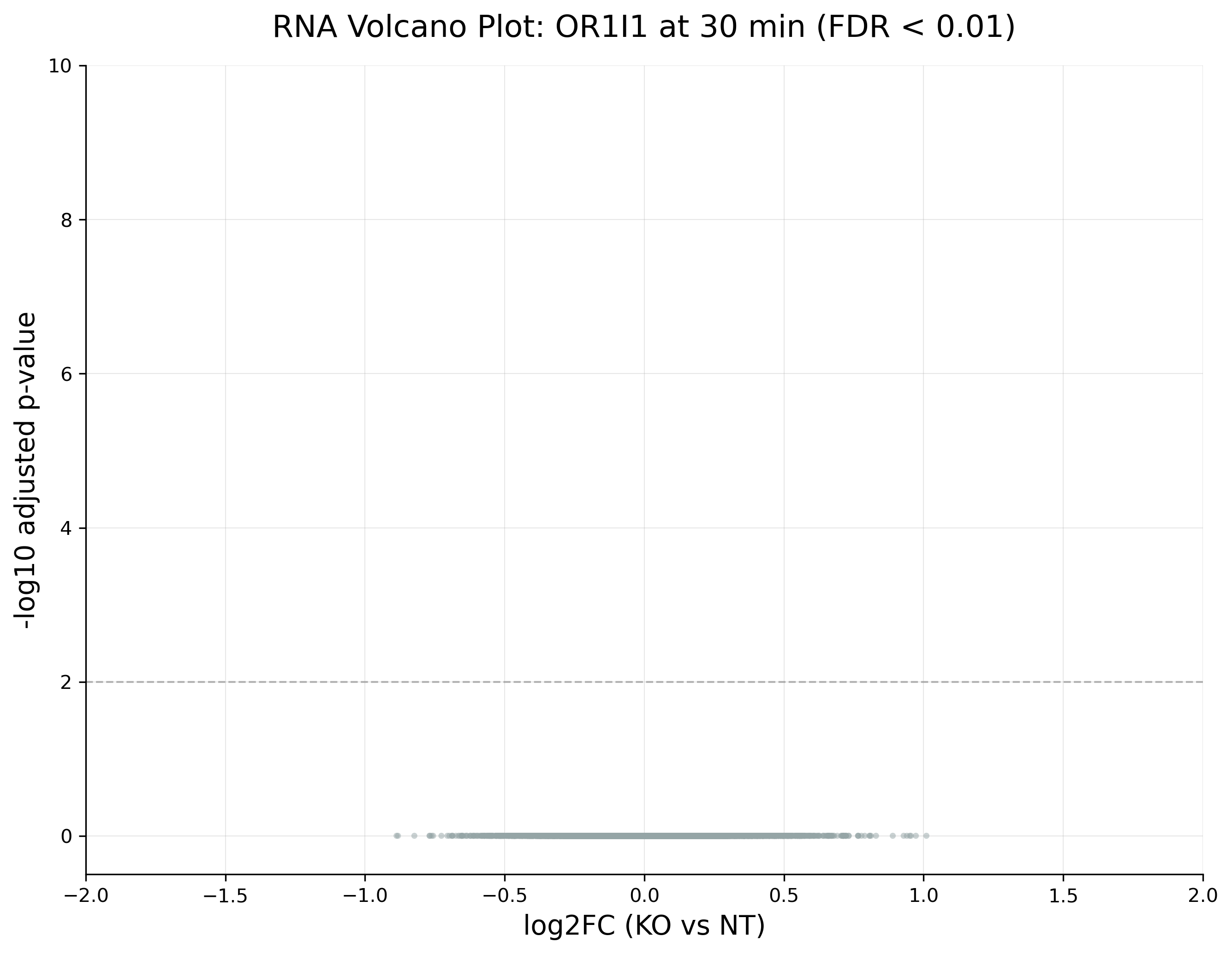

Supplementary figure 6: Volcano plot of differential RNA abundance (OR1I1-KO vs NT) at 30 min IL-1β treatment.
